# A binary trait model reveals the fitness effects of HIV-1 escape from T cell responses

**DOI:** 10.1101/2024.03.03.583183

**Authors:** Yirui Gao, John P. Barton

**Affiliations:** Department of Physics and Astronomy, University of California, Riverside, USA; Department of Computational and Systems Biology, University of Pittsburgh School of Medicine, USA; Department of Physics and Astronomy, University of Pittsburgh, USA

## Abstract

Natural selection often acts on multiple traits simultaneously. For example, the virus HIV-1 faces pressure to evade host immunity while also preserving replicative fitness. While past work has studied selection during HIV-1 evolution, as in other examples where selection acts on multiple traits, it is challenging to quantitatively separate different contributions to fitness. This task is made more difficult because a single mutation can affect both immune escape and replication. Here, we develop an evolutionary model that disentangles the effects of escaping CD8^+^ T cell-mediated immunity, which we model as a binary trait, from other contributions to fitness. After validation in simulations, we applied this model to study within-host HIV-1 evolution in a clinical data set. We observed strong selection for immune escape, sometimes greatly exceeding past estimates, especially early in infection. Conservative estimates suggest that roughly half of HIV-1 fitness gains during the first months to years of infection can be attributed to T cell escape. Our approach is not limited to HIV-1 or viruses, and could be adapted to study the evolution of quantitative traits in other contexts.

## Introduction

Natural selection acts on phenotypic traits, as individuals with traits that are well-adapted to their environment enjoy relative reproductive success. Understanding how selective pressures shape evolution is a major goal of evolutionary biology. Applications include studying the drug resistance in bacteria ^1^ and patterns of evolution in cancer ^2^ or viruses ^3^.

In population genetics, methods have been developed to detect natural selection by examining the genetic diversity of a population and how it changes over time ^4–11^. However, inferred selection for a particular allele is not always easy to interpret. A single mutation can affect multiple traits, each of which may be subject to selection. Many phenotypic traits are also polygenic, meaning that they are affected by genetic variation at multiple sites and/or genes ^12,13^. How can selection for a particular trait be disentangled from other evolutionary forces?

To approach this question, we developed a model to quantify selection on a binary trait, distinct from other contributions to fitness, using sequence data from an evolving population. Our model is motivated in particular by human immunodeficiency virus (HIV-1) infection. HIV-1 evolves rapidly during infection, and specific mutations allow the virus to escape immune control ^14–17^. Past work has shown that the pressure to escape from CD8^+^ T cell responses is particularly strong ^11,18–20^, though this is difficult to quantify precisely ^20–22^. While CD8^+^ T cell escape mutations can rapidly sweep through viral populations, the same mutations also often revert when the virus is transmitted to a new individual with a different immune response ^18,20,23–25^. This suggests that some escape mutations may harm viral replication in the absence of immune pressure.

After testing our approach in simulations, we applied it to study viral evolution in a data set derived from 13 HIV-1-infected individuals. In this data, we observed strong selection for CD8^+^ T cell escape independent of other contributions to fitness. This was balanced by a decrease in the estimated fitness effects of specific escape mutations, compared to estimates in a model without the escape trait. Remarkably, we found that fitness gains due to CD8^+^ T cell escape were responsible for roughly half of the gains in viral fitness estimated from data, especially early in infection. Reversions to clade consensus were also strongly selected, comprising roughly a quarter of viral fitness gains after several years of evolution. Thus, in this data set we found that the vast majority of HIV-1 adaptation was driven by CD8^+^ T cell escape and reversions.

While the current work focuses on HIV-1 evolution, our framework for estimating the strength of selection on a binary trait is generic. This approach could be applied to study trait evolution in other contexts using temporal sequence data.

## Results

### CD8^+^ T cell escape as a binary trait

T cells are a vital part of the adaptive immune system. A subset of T cells that express CD8 on the cell surface specializes in eliminating intracellular pathogens and cancer. Each T cell is equipped with a T cell receptor (TCR) that binds to a specific peptide, roughly 10 amino acids in length, which can be displayed by major histocompatibility complex type I (MHC-I) molecules on the target cell surface. When an activated CD8^+^ T cell binds to its cognate antigen, this can trigger the release of cytotoxins that kill the target cell. CD8^+^ T cells that recognize peptides derived from viral proteins can thus play an important role in controlling infection by killing infected cells before they have the chance to release viral particles. The importance of cytotoxic T cells in the context of HIV-1 infection has been appreciated since the early days of HIV-1 research ^26^.

Here we model the susceptibility of HIV-1 viruses to killing by CD8^+^ T cell clones as a binary trait. That is, for each HIV-1-specific T cell clone within an individual, each virus is either vulnerable (0) or not (1). This is determined by whether or not viral proteins contain the peptide that a T cell binds, which is also referred to as an epitope.

In principle, one may imagine that states of partial immunity are possible: a virus may not express precisely the same epitope that a CD8^+^ T cell recognizes, but it may express one that is similar enough that it can also be detected by T cells. However, the binding of TCRs to peptide-MHC-I complexes is highly specific. While experiments have shown that some TCRs are cross-reactive to highly similar peptides, most mutations within the epitope abrogate T cell recognition ^27–30^. Thus, as we describe below, we modeled HIV-1 susceptibility to CD8^+^ T cells as “all or nothing,” where any nonsynonymous mutation within a T cell epitope results in a loss of immune recognition.

### Evolutionary model including selection on binary traits

Our starting point is a Wright-Fisher model, which describes the evolution of a population of *N* individuals represented by genetic sequences of length *L*. For simplicity, in the main text we will assume that genetic variants are binary, with each locus consisting of either a wild-type or mutant allele (see Methods for a more detailed model). We write the fitness *f*_*a*_ of an individual with sequence *a* as

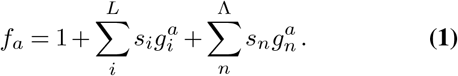

Here, the *s*_*i*_ are selection coefficients that quantify the fitness effects of the mutant allele at each site *i*. The 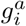 are indicator variables, with 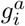 equal to one if sequence *a* has a mutant allele at site *i*, and zero otherwise. The last terms in (1) give the fitness contributions of the binary traits (CD8^+^ T cell escape, in our case; see **Fig. 1e** for an example). The *n* index labels different CD8^+^ T cell epitopes, with *s*_*n*_ quantifying the fitness effect of escape for the epitope labeled *n*. Unlike a regular locus, the binary trait can be seen as a “virtual” locus. Since multiple mutations within an epitope can confer escape, determining *g*_*n*_ requires considering genetic variation at multiple loci. We set 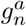 to one if sequence *a* contains nonsynonymous mutations within epitope *n*, and zero otherwise. This model extends the simple additive fitness model used in a prior study of HIV-1 evolution, where the fitness benefits and costs of escape were effectively combined together into the selection coefficients for escape mutations ^11^. During each generation of replication, the probability that an individual replicates is proportional to its fitness.

**Fig. 1.**
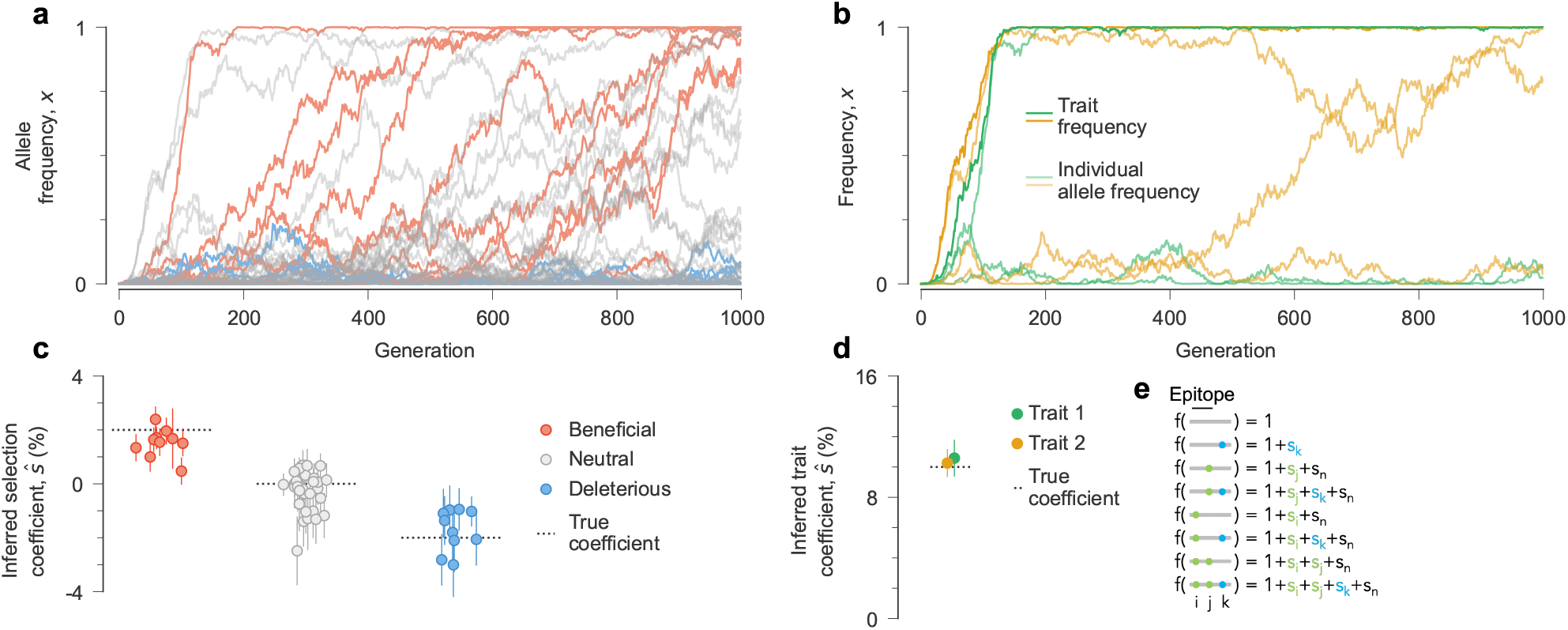
Recovering selection on individual mutations and traits from temporal genetic data. **a**, Simulated mutant allele frequency trajectories. **b**, Trait frequencies and their contributing individual mutant allele frequencies in the same simulation. Trait frequencies can approach one even when the frequencies of contributing alleles remain modest. The fitness contributions of individual mutations (**c**) and traits (**d**) that we infer are close to their true values. **e**, Illustration of the fitness function (1). Nonsynonymous mutations within an epitope affect both escape and intrinsic fitness through distinct terms. We assume that a single nonsynonymous mutation within an epitope is sufficient for escape. Simulation parameters: *L* = 50 loci with two alleles at each locus (mutant and WT), ten beneficial mutants with *s* = 0.02, 30 neutral mutants with *s* = 0 and ten deleterious mutants with *s* = −0.02. We consider two binary traits, each with three contributing alleles and trait coefficients *s* = 0.1. Mutation probability per site per generation *µ* = 2 *×* 10^−4^, recombination probability per site per generation *r* = 2 *×* 10^−4^, population size *N* = 10^3^. The initial population contains all WT sequences, evolved over *T* = 1000 generations.

In addition to natural selection, our model includes spontaneous mutations, recombination, and genetic drift. We assume a simple probability *µ* per site per generation for the allele at each site *i* to change from wild-type (WT) to mutant (and vice versa). This can easily be extended to model different, asymmetric mutation rates between nucleotides in a realistic sequence model (Methods). Genetic drift due to the finite population size of *N* individuals adds randomness to the evolutionary dynamics.

For HIV-1, recombination can occur when two different viruses coinfect the same cell and RNA from each of them is packaged in new virions. When virions containing distinct RNAs infect new cells, the reverse transcriptase can switch between RNA templates, producing recombinant DNA. The two components of this process are the coinfection rate, *p*_*c*_, and the template switching rate, *p*_*s*_. Estimates of the effective recombination rate *r* ∼ *p*_*c*_*p*_*s*_ are high, with *r* = 1.4 × 10^−5^ (ref. ^31^). This is comparable to estimates of HIV-1 mutation rates ^32^. Following recent work showing that HIV-1 recombination occurs more frequently in individuals with higher viral loads due to higher coinfection probabilities ^33^, we allowed the overall recombination rate to vary over time along with the measured viral load within each individual (Methods).

### Inference of natural selection from temporal genetic data

We developed a method to infer selection, including both selection coefficients *s*_*i*_ for individual mutations and coefficients *s*_*n*_ for binary traits, from sequence data sampled over time. Our approach works by computing the probabilities of different evolutionary histories (i.e., frequencies of individual sequences in the population over time) as a function of the selection coefficients. We then find the selection coefficients that maximize the posterior probability of the data using Bayes’ theorem.

To make this inference problem tractable, we used a number of approximations and analytical techniques. First, we considered our evolutionary model in the so-called diffusion limit ^34^. In this limit, we assume that the population size *N* is very large, and that the selection coefficients *s*_*i*_, trait coefficients *s*_*n*_, mutation probabilities *µ*, and recombination probability *r* are small. We can then derive effective equations for the dynamics of the allele frequencies *x*_*i*_ and the escape trait frequencies *x*_*n*_, which are described by a Fokker-Planck equation (Methods). Allele frequencies *x*_*i*_ are defined as the fraction of individuals in the population with a mutant allele at site *i*, and trait frequencies *x*_*n*_ represent the fraction of individuals with a mutant allele at least one of the epitope sites for epitope *n*.

While the effective equations are mathematically complicated, their meaning can be clearly interpreted. Natural selection, mutation, and recombination drive deterministic changes in allele and/or trait frequencies, while the finite population size contributes to noise. Alleles and traits are likely to increase in frequency if they appear on sequences that have higher fitness than the average fitness of the population. Mutation introduces new genetic variation, driving allele and trait frequencies away from zero or one. The effects of recombination are more subtle, with no net effect on expected changes in allele frequencies. However, recombination can drive changes in trait frequencies in a way that depends on the arrangement of mutations within the epitope (see Methods).

Finally, we used methods from statistical physics to convert the Fokker-Planck equation into a path integral ^11,35–38^ (Methods). The path integral quantifies the likelihood of different evolutionary histories or “paths” as a function of the selection coefficients. We can then derive an analytical expression for the vector of selection coefficients ***ŝ***, including both the *s*_*i*_ and the *s*_*n*_, that best fits the data (see Methods),

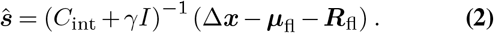

Here *C*_int_ is the allele/trait frequency covariance matrix integrated over time, which accounts for the speed of evolution and the correlations between mutations/traits. The parameter *γ* specifies the width of a Gaussian prior distribution for the selection coefficients, and *I* is the identity matrix. The net change in allele/trait frequencies over the trajectory is Δ***x***. Finally, ***µ***_fl_ and ***R***_fl_ quantify the expected cumulative change (or flux) in allele/trait frequencies due to mutation and recombination, respectively. Intuitively, (2) attributes net changes in allele/trait frequencies that are not already explained by mutation or recombination to natural selection. The speed of evolution and the genetic background then control how beneficial or deleterious an allele or trait is inferred to be.

### Performance on simulated data

To benchmark the performance of our method, we simulated population evolution in the Wright-Fisher model using the fitness function defined in (1) (**Fig. 1**). We considered sequences with *L* = 50 loci (10 beneficial with selection coefficients *s* = 2%, 30 neutral, and 10 deleterious with selection coefficients *s* = −2%), with a population size of *N* = 1000 individuals. We also included strong selection (*s* = 10%) on Λ = 2 binary traits, each with three contributing alleles that were randomly chosen across the sequence. Individuals thus receive a substantial increase in fitness for mutations at at least one of the trait sites, but multiple mutations confer no additional benefit. **Figure 1** shows that our method is able to distinguish between beneficial, neutral, and deleterious alleles, and to estimate selection on these binary traits. When selection varies over time, the constant coefficients that we infer are typically similar to time-averaged ones ^11^ (**Supplementary Fig. 1**).

### Robustness to limited sampling

Real data sets often face significant limitations in both the number of sequences that can be obtained and the frequency of sampling. To test the robustness of our approach to finite sampling, we ran 100 simulations across a wide range of sampling conditions. We assessed our ability to accurately identify selection by computing the average area under the receiver operating characteristic (AUROC) for classifying beneficial/deleterious mutations. We also computed the normalized root mean square error (NRMSE) for the inferred trait coefficients. While inference becomes more difficult with limited sampling, declines in performance occur gradually (**Supplementary Fig. 2**). Our method is particularly robust to limited numbers of sequences, with the ability to identify beneficial or deleterious mutations using as little as 10 sequences per time point.

### Selection for immune escape in intrahost HIV-1 evolution

We applied our approach to study HIV-1 evolution using 13 patient data sets ^39^ (Methods). For each individual, longitudinal HIV-1 half-genome sequences were collected from around the time of peak infection up to several months or years afterward. This time spans both the acute and chronic phases of infection. Early-phase CD8^+^ T cell epitopes were also carefully verified ^30,39^, which allowed us to identify putative CD8^+^ T cell escape mutations. We then estimated the selection coefficients for individual mutations and T cell escape across all 13 individuals. Donors did not receive antiretroviral drug treatment during this study.

In order to confidently separate the fitness contribution of escape as a trait from the individual escape mutations, we focused on a well-sampled subset of all T cell epitopes. Specifically, we estimated escape coefficients for epitopes where at least three putative escape mutations were observed. Out of 71 CD8^+^ T cell epitopes in this data set, 38 had at least three escape mutations, thus allowing us to estimate their escape coefficients. For the remaining 33 epitopes, we still estimated selection coefficients for individual escape mutations, but we did not attempt to disentangle the contributions of T cell escape versus intrinsic fitness.

Overall, we found very strong selection for CD8^+^ T cell escape in most epitopes (**Fig. 2a**). Inferred escape coefficients ranged from nearly neutral to highly beneficial (*s* ∼ 26%), with a mean value of 7%. **Figure 3** shows a typical example of HIV-1 evolution to escape from T cell responses across three epitopes in individual CH470. Here, multiple escape mutations are observed for each epitope, making it challenging to infer the strength of selection for escape by looking at single mutations alone.

**Fig. 2.**
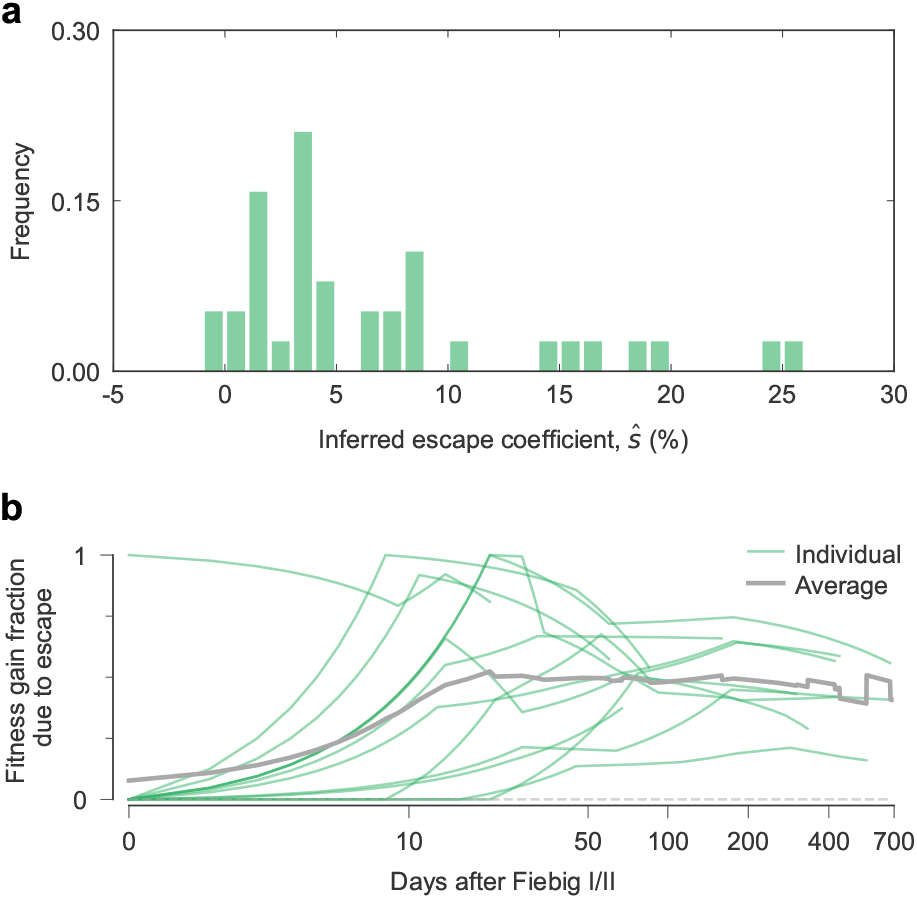
CD8^+^ T cell escape is strongly selected and contributes substantially to intrahost HIV-1 fitness gains. **a**, Distribution of inferred fitness effects of CD8^+^ T cell escape in HIV-1 patient data. Mutations within T cell epitopes allow infected cells to escape killing by T cells, thus allowing the virus to continue productive replication. **b**, The fraction of the total increase in within-host HIV-1 fitness inferred for each of 13 individuals that is due to CD8^+^ T cell escape.

**Fig. 3.**
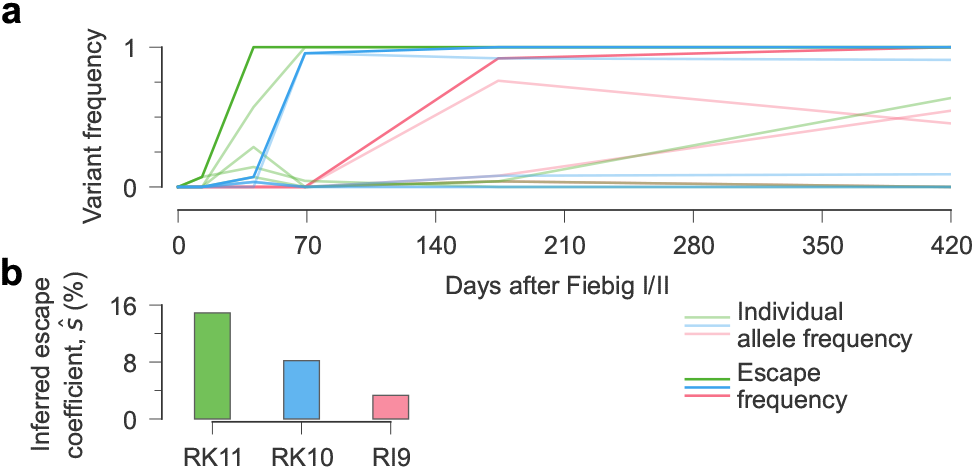
Estimates of selection for CD8^+^ T cell escape in a simple example. **a**, Frequency of individual escape mutations and the escape trait for three epitopes in the 3^*′*^ half-genome for individual CH470. Escape in each epitope is mediated by multiple mutations. **b**, We infer strong positive selection for escape for all three epitopes.

Next, we quantified how much CD8^+^ T cell escape contributes to observed changes in HIV-1 fitness *in vivo*. To do this, we computed the fitness of each sequence in each individual and measured the fraction of fitness increase (relative to the transmitted/founder (TF) virus) due to escape coefficients. Here we also included the contribution from escape mutations in epitopes where escape coefficients were not separately estimated. We observed a consistent pattern where escape mutations were rapidly selected within the first weeks of infection, followed by a long, slow decline (**Fig. 2b**). In only 10 days, T cell escape comprises roughly half of HIV-1 fitness gains. This is especially remarkable given that the identified T cell epitopes cover less than 2% of the HIV-1 proteome in these individuals.

### Reversions and the intrinsic fitness effects of CD8^+^ T cell escape mutations

CD8^+^ T cell escape mutations have been observed to revert after the virus is transmitted to a new host ^18,20,23–25^, suggesting that escape mutations may be deleterious in the absence of immune pressure. Before explicitly modeling T cell escape as a trait, a large fraction (35%) of escape mutations are inferred to be substantially beneficial (*ŝ >* 1%; **Supplementary Fig. 3**). When escape coefficients are included, the distribution of inferred selection coefficients for escape mutations shifts substantially towards more deleterious values, with a peak near zero. In this case, only 15% of escape mutations are inferred to be substantially beneficial. HIV-1 evolutionary dynamics within each host are therefore best explained by a model where CD8^+^ T cell escape provides a substantial fitness benefit, which is achieved through escape mutations that may be nearly neutral or even moderately deleterious.

**Figure 4** shows an example of the shift in inferred selection coefficients for escape mutations after escape is modeled as a trait. Escape at the EV11 epitope targeted by individual CH131 occurs rapidly. However, there is no single, primary escape mutation. Instead, a set of six escape mutations compete for dominance in the viral population. Without the inclusion of an escape coefficient, the inferred selection coefficients for all of these mutations are positive, with a maximum of around 6%. When we estimate an escape coefficient for EV11, the inferred selection coefficients for individual escape mutations shift to appear more deleterious. In addition, the inferred fitness benefit of escape for the EV11 epitope is around 15%, far higher than the selection coefficient for any individual escape mutation that we estimated in the absence of the escape coefficient.

**Fig. 4.**
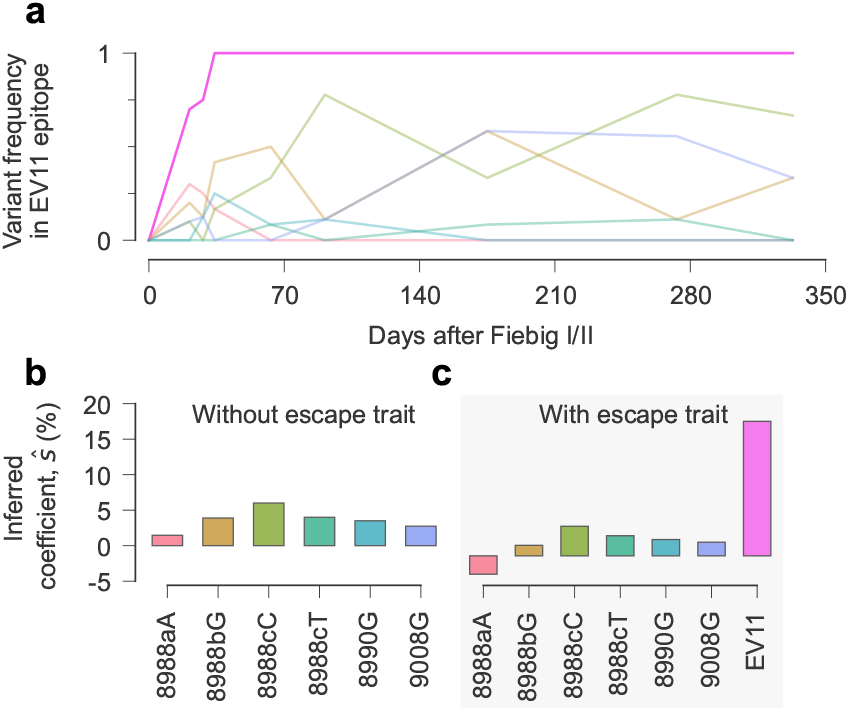
Inclusion of escape as a trait increases the inferred selective advantage of escape and decreases the apparent advantage of individual escape mutations. **a**, Allele frequencies and escape frequency for epitope EV11 in individual CH131. Escape at this epitope is complex, with contributions from six distinct mutations. **b**, Without treating escape as a trait, all EV11 escape mutations are inferred to be positively selected. **c**, When the selective advantage of escape is considered separately, the inferred selection coefficients for individual mutations become more deleterious. Conversely, the estimated advantage of EV11 escape is substantial, much larger than the inferred selection coefficient for any individual escape mutation without the escape trait.

The intrinsic fitness cost of escape mutations is also high-lighted by the benefit of reversions toward the clade consensus sequence. Similar to our previous analysis for CD8^+^ T cell escape mutations, we computed the fraction of HIV-1 fitness gains in each individual that can be attributed to reversions over time. We observed that reversions contributed substantially to HIV-1 fitness *in vivo*, in a way that grows steadily over time (**Supplementary Fig. 4**). After several years of infection, an average of around 25% of HIV-1 fitness gains relative to the TF virus are due to reversions.

For both T cell escape and reversions, we note that fitness contributions are heterogeneously distributed across individual mutations. Inferred selection coefficients for both types of mutations have long tails (**Supplementary Fig. 5**). A few, large selection coefficients thus make large contributions to fitness.

### Stability of inferred selection outside T cell epitopes

Changing the fitness model that we use to describe HIV-1 evolution can affect the fitness effects of mutations that we infer. For mutations within CD8^+^ T cell epitopes, these effects can be quite large, as described above (see also **Supplementary Fig. 3**). Does the inclusion of escape coefficients also affect mutations outside of T cell epitopes? Large changes in these selection coefficients could indicate that the inference process is highly sensitive to specific modeling choices, casting doubt on its reliability.

Overall, the inclusion of escape coefficients shifts the inferred effects of escape mutations, which are typically inferred to be highly beneficial without considering the escape trait, toward much more deleterious values, but without substantially reshaping the overall distribution of selection coefficients (**Supplementary Fig. 5**). **Figure 5** shows a representative example, demonstrating that inferred selection coefficients for individual mutations with and without the inclusion of escape coefficients are very strongly correlated. Most selection coefficients that change significantly are located within CD8^+^ T cell epitopes, as expected. However, the selection coefficient for one non-epitope mutation, 974A, undergoes a substantial change (*ŝ*∼ 3% to around 2% before and after the inclusion of escape coefficients, respectively). We found that 974A was strongly linked with 3951C, an escape mutation in the RL9 epitope, which potentially explains this difference.

**Fig. 5.**
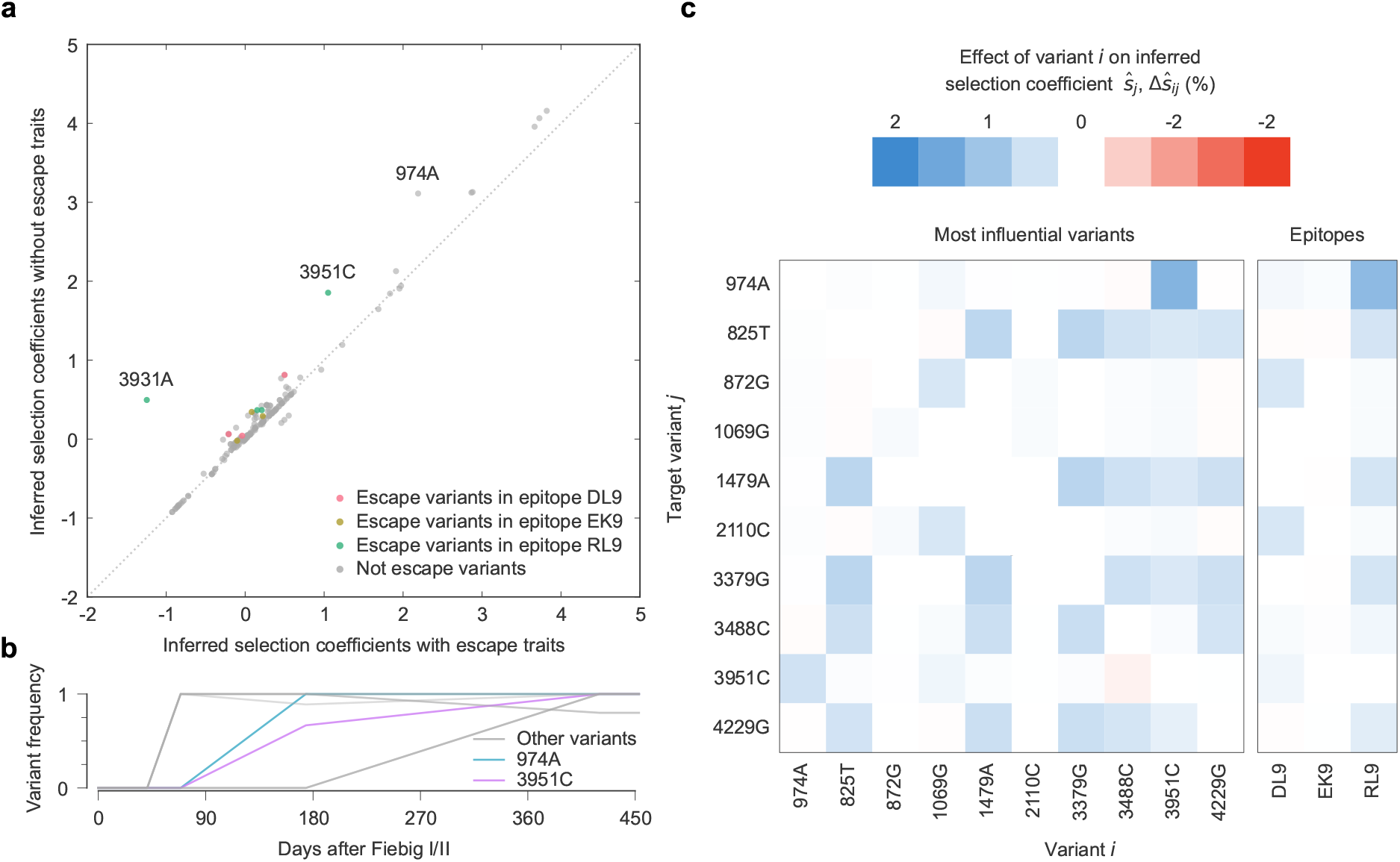
Inclusion of escape as a trait only strongly affects inferred selection coefficients for escape mutations and alleles that are strongly linked to them. **a**, Inferred selection coefficients with and without escape terms inferred from 5^*′*^ HIV-1 half-genomes from individual CH470. There are three alleles with substantial shifts in inferred selection. Two of these (3931A and 3951C) are escape variants in RL9. **b**, Allele frequencies for variants that appear frequently with 974A, demonstrating strong linkage with 3951C. **c**, Linkage effects on inferred selection coefficients for mutations linked to 974A, and escape coefficients for the three CD8^+^ T cell epitopes. Effects shown the strong interaction between variant 974A and variant 3951C, with RL9 escape decreasing the inferred selection coefficient for 974A.

In this example, we comprehensively explored the effects of linkage disequilibrium (i.e., correlations) between mutations/escape on inferred selection coefficients. Estimated selection coefficients for all mutations and traits are connected due to the integrated allele/trait frequency covariance matrix *C*_int_ in (2). For each escape mutation and trait *j*, we computed the selection coefficients that would be inferred if each other mutation/trait *i* was ignored, denoted 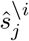 (see Methods). The difference between the coefficient inferred for *j* using the full data and the coefficient inferred when mutation/trait *i* is held out yields the effect of linkage with *i* on *j* (ref. ^11^),

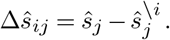

**Figure 5c** shows Δ*ŝ*_*ij*_ values for 974A, the escape coefficients for epitopes DL9, EK9, and RL9, and the individual escape mutations that occur within these epitopes. Here we find that linkage disequilibrium with 3951C and RL9 escape mutants is indeed primarily responsible for the shift in selection inferred for 974A. The fitness benefit of escape partly explains the rise to fixation of 974A, reducing its estimated selection coefficient.

## Discussion

In this paper, we developed a new method to estimate the fitness effect of a binary trait, jointly with selection coefficients for individual mutations, from temporal genetic data. After validation in simulations, we applied this method to study intrahost HIV-1 evolution in a clinical data set. We modeled CD8^+^ T cell escape as a binary trait to disentangle the contributions of immune evasion and intrinsic replication to HIV-1 fitness. In this data, HIV-1 evolution was best explained by models with strong selection for T cell escape, but with individual escape mutations that are typically nearly neutral. Reversions to clade consensus were also inferred to be beneficial. The contributions of reversions to HIV-1 fitness gains grew steadily over time.

Our findings are consistent with past observations that have reported strong selection for T cell escape ^11,18,20,39–41^. However, here we found an even stronger benefit for CD8^+^ T cell escape compared to past work ^11^. There are also reasons to believe that the selection for T cell escape may be even stronger than what we have estimated. The Bayesian regularization that we have used controls the inference of large selection coefficients that are not well-supported by data, but it favors spreading the effects of a single large parameter (e.g., a large escape coefficient) into multiple smaller ones (e.g., selection coefficients for individual escape mutations) to maximize the posterior distribution. This is a well-known effect of *L*_2_-norm regularization. This effect could be alleviated in future work through the choice of a different prior distribution, though this could make the analytical derivation of the estimator for selection/trait coefficients more challenging.

The apparent selective advantage of reversions has also been noted in prior work ^11,20,25^. However, to our knowledge, the relative genome-wide contributions of immune escape across CD8^+^ T cell epitopes and reversions to HIV-1 fitness, and how these contributions vary over the first few years of infection, have not previously been quantified. Remarkably, after several years of infection, roughly three quarters of within-host HIV-1 fitness gains can be attributed to CD8^+^ T cell escape or reversions (**Supplementary Fig. 6**). As discussed by Zanini and collaborators ^20^, the apparent consistency of selective pressures on HIV-1 across many infected individuals, reflected by strong selection for reversions, could help to explain the success of HIV-1 fitness models that are based on sequence conservation across hosts ^30,42–44^.

Several previous studies have attempted to separately quantify the fitness benefits and costs of T cell escape and escape mutations, respectively, in HIV-1 or SIV (refs. ^40,41,45,46^). In contrast with the present work, these studies fit escape benefits and fitness costs by comparing two populations of individuals: one in which a particular T cell epitope is targeted, and another (often HLA-mismatched) one that does not respond in the same way. Estimates of the fitness effects of escape vary. Some studies found very strong effects for particular epitopes ^40,41^, whereas others estimated both escape benefits and costs that are very close to zero ^46^. Compared to these studies, the typical fitness benefit of escape that we observe is more moderate, with a median selection coefficient of around 4% (similar to ref. ^45^). However, as shown in **Fig. 2**, the fitness benefit of escape for some epitopes is inferred to be as high as 25%. Our results are most different from those of ref. ^46^, which may be partly attributable to differences in data sets. In the data set that we use, viral sequences were obtained frequently, especially early in infection, and donors were not using antiretroviral drugs. For ref. ^46^, the shortest time between longitudinal samples was around 6 months, and many individuals received some drug treatment. Our work also accounts for interference between viral variants, which can lead to much larger estimates of the fitness effects of beneficial mutations ^11^. While the inferred fitness effects of mutations for genetically similar viruses can be remarkably well-correlated across different hosts ^47^, some differences between our results and prior studies may also be due to the joint estimates of fitness effects across multiple individuals in prior work.

Our study has some limitations due to the constraints on the data available. While our approach is very robust to limited numbers of sequences, long gaps in time between samples limit our ability to fully resolve evolutionary dynamics, leading to less accurate inferences (**Supplementary Fig. 2**). In T cell epitopes, when there are few observed escape mutations it is also statistically challenging to distinguish between the fitness effects of individual mutations and the fitness benefit of escape. This is why we have not attempted to estimate escape coefficients for epitopes with only one or two escape mutations.

The fitness model that we have used could also be extended. While the form of our model was motivated by the study of HIV-1 escape from CD8^+^ T cell responses, it could apply binary traits in other evolutionary contexts as well. In future work, we aim to extend this approach to estimate selection on more general quantitative traits. Ultimately, the development of new methods to fit more realistic models to data will improve our quantitative understanding of evolution.

Future studies should also consider time-varying selection. In principle, we may expect to observe time-varying selection for T cell escape as immune responses rise and fall. With our current approach, the selection coefficients that we estimate are roughly equivalent to the average selection coefficient when selection changes in time ^11,37^ (see **Supplementary Fig. 1**). In epitopes that accumulate multiple escape mutations, however, the trait coefficients that we estimate may be more similar to the strength of selection that originally drove escape, at least in cases where the individual escape mutations are not strongly deleterious. This is because the rate for escape to “revert” can be quite slow when most viruses possess multiple escape mutations, requiring more than one mutation or recombination event for all escape mutations to be removed. In this case, it could be difficult to accumulate evidence for weaker selection for escape because sequences without any escape mutations are only produced rarely. Future work should examine how selection for immune escape varies with time, and how this is associated with the corresponding strength of immune responses.

## ACKNOWLEDGEMENTS

The work of Y.G. and J.P.B. reported in this publication was supported by the National Institute of General Medical Sciences of the National Institutes of Health under Award Number R35GM138233.

## AUTHOR CONTRIBUTIONS

All authors contributed to methods development, data analysis, interpretation of results, and writing the paper. J.P.B. supervised the project.

## Methods

### Evolutionary model with selection on binary traits

In our study, model the evolution of a population of *N* individuals subject to recombination, mutation, and natural selection, following the Wright-Fisher (WF) model. For simplicity, we first describe a model with only two alleles per site, wild-type (WT) or mutant (MT). Individuals possess a genetic sequence length of *L*, resulting in a total of *M* = 2^*L*^ genotypes. We further assume that there exist Λ binary traits, which depend on the presence or absence of mutant alleles at specific sites, and which are also subject to selection. For clarity, we will use *i, j*,… indices to denote the different loci or sites in the sequence, *n, m*, … are used for trait indices, and *a, b*, … represent genotype indices. Finally, *t*_1_, *t*_2_, …, *t*_*K*_ are used as generation (time) indices. The first two sets of indices are presented as subscripts, the genotype indices as superscripts, and the generation indices are displayed within parentheses to indicate quantities that vary with time.

Let *n*_*a*_(*t*_*k*_) represent the number of individuals with genotype *a* at generation *t*_*k*_, with *z*_*a*_(*t*_*k*_) = *n*_*a*_(*t*_*k*_)*/N* the frequency of genotype *a* at generation *t*_*k*_. The vector *z*(*t*_*k*_) = (*z*_*a*_(*t*_*k*_), *z*_*b*_(*t*_*k*_), …, *z*_*M*_ (*t*_*k*_)) describes the state of the population at generation *t*_*k*_. In our model, the probability of recombination occurring per site per generation is denoted by *r*. After recombination, the mean frequency of genotype *a* at generation *t*_*k*+1_, is

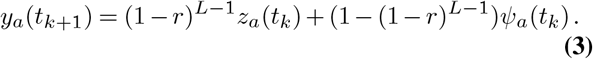

The term (1 − *r*)^*L*−1^ gives the likelihood of an individual not experiencing recombination, and *ψ*_*a*_(*t*_*k*_) denotes the probability that the random recombination of any two individuals within the population results in an offspring of genotype *a*. This includes scenarios where both parent and offspring share the same genotype *a*.

After recombination, the mean frequency of each genotype in the next generation *p*_*a*_(*t*_*k*_) is shaped by selection and mutation,

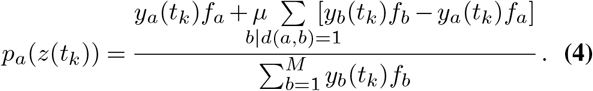

In this case, all alleles and loci share an equal mutation rate *µ*. For simplicity, we assume that each sequence undergoes at most one mutation per generation, given the exceedingly low mutation rate *µ*. The notation *b*|*d*(*a, b*) = 1 indicates that genotypes *a* and *b* differ by just a single mutation.

Due to the highly specific nature of TCR binding to peptide-MHC-I complexes, most nonsynonymous mutations within an epitope are likely to significantly disrupt binding, thereby facilitating immune escape. For any given epitope, different mutation paths can lead to a similar outcome: loss of recognition by T cells. Therefore, T cell escape for each individual epitope can be effectively modeled as a binary trait. The fitness of genotype *a*, denoted as *f*_*a*_, is given by

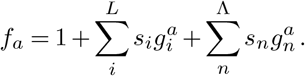

Contributions to fitness come from the effects of individual alleles (quantified by *selection coefficients s*_*i*_) and traits (*trait coefficients s*_*n*_). For the former, the impact of mutations is cumulative. In other words, the effects of mutant alleles at different sites add together. In contrast, for any given epitope *n*, the presence of one or more nonsynonymous mutations results in *g*_*n*_ being assigned a value of 1, irrespective of the number of these mutations. However, we emphasize that the fitness effects of different trait terms are additive, so that the effects of escape in two *different* T cell epitopes will add.

### Path integral likelihood

Under WF dynamics, the probability of observing genotype frequencies *z*(*t*_*k*+1_) at generation *t*_*k*+1_, given genotype frequencies of *z*(*t*_*k*_)= (*z*_*a*_(*t*_*k*_), *z*_*b*_(*t*_*k*_), …, *z*_*M*_ (*t*_*k*_)) at generation *t*_*k*_, is

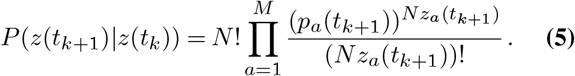

Consequently, the likelihood that the genotype frequency vector follows a specific evolutionary trajectory, or “path”, ***z*** = (*z*(*t*_1_), *z*(*t*_2_), · · ·, *z*(*t*_*K*_), is

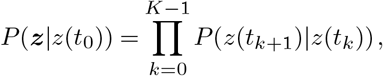

conditioned upon the initial state *z*(*t*_0_). This expression is difficult to work with directly, so we use several approximations to make our analysis more tractable.

To simplify this expression, we will project dynamics to the level of individual mutant loci *i* and trait groups *n*, instead of genotypes *a*. Here, we use the term “trait group” to refer to the set of all loci that contribute to the same CD8^+^ T cell epitope. We denote the frequency of mutant alleles at site *i* in the population as *x*_*i*_, and the frequency of individuals with one or more nonsynonymous mutant alleles in trait group *n* as *x*_*n*_. Additionally, the frequency of paired mutant alleles *x*_*ij*_ is used to describe the correlation between different mutations. The formulas for these are as follows:

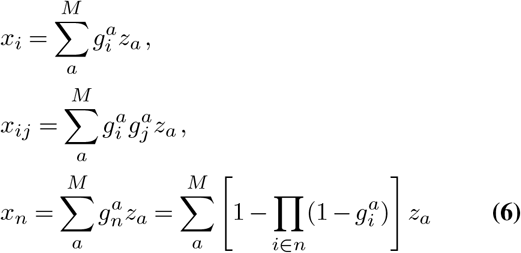

Here the term 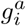 indicates whether genotype *a* contains a mutant allele at locus *i*, with the wild type (WT) having 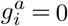 and the mutant type (MT) having 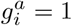. Similarly, 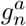 specifies whether trait group *n* contains a mutant allele. If all loci in trait group *n* of genotype *a* are WT, then 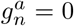; however, if there is at least one mutant allele in the trait group, meaning that at least one locus within the trait group has 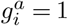, then 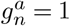.

It is worth noting that, in most cases, trait groups can be regarded as a special type of locus. Consequently, we utilize the new subscripts ***i, j*** to represent generic loci — encompassing both individual loci and trait groups — to distinguish them from individual loci *i, j*. Taking the pair allele frequency as an example, *x*_***ij***_ will represent not only the correlation between different individual loci but also between an individual locus and a trait group, as well as between different trait groups.

Next we consider the dynamics of the mutant allele frequencies (and trait frequencies) in the diffusion limit ^48^. We assume that the population size *N* → ∞ and that the selection coefficients, mutation rate, and recombination rate are all small (𝒪(1*/N*)). In this limit, applying methods from statistical physics, the probability of an evolutionary trajectory ***x*** = (*x*(*t*_1_), *x*(*t*_2_), …, *x*(*t*_*K*_)) can be quantified using a path integral (see refs. ^49–51^ for more details on this approach)

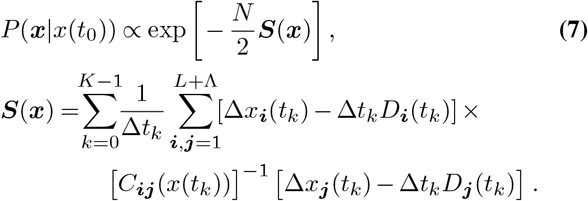

Here Δ*t*_*k*_ = *t*_*k*+1_ − *t*_*k*_ and Δ*x*_***i***_(*t*_*k*_) = *x*_***i***_(*t*_*k*+1_) − *x*_***i***_(*t*_*k*_). ***S***(***x***(*t*_*k*_)) is referred to as the action in physics. In this expression, trait terms are considered as special individual loci, thus the total length of frequencies is the length of generic loci, which is *L* + Λ (binary case). In statistical physics, *D*_***i***_(*t*_*k*_) and *C*_***ij***_(*t*_*k*_)*/N* are commonly referred to as the drift vector and the diffusion matrix respectively, which will be discussed in the following section. To prevent confusion, here we note that the drift vector quantifies the effects of natural selection, mutation, and recombination, which affect the expected change in allele frequencies, and *not* genetic drift.

### The drift vector and diffusion matrix

In this section, unless otherwise specified, the time parameter for all physical quantities is assumed to be *t*_*k*_, and the corresponding time indices will be omitted. We first begin with the diffusion matrix *C*_***ij***_*/N*, which is the same for both allele and trait frequencies

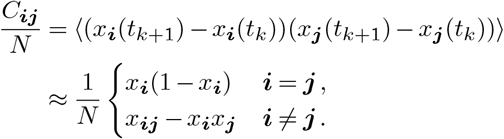

The expectation in the first line is taken over the WF model dynamics, which yields the second line in the limit that *N* is large and terms of 𝒪 (1*/N* ^2^) are omitted. Here ***i, j*** can be used in both individual and trait terms. *x*_***i***_ is the mutant frequencies at generic loci ***i***, while *x*_***ij***_ is the frequency of individuals with mutations at both generic loci ***i*** and ***j*** (see (6)). The diffusion matrix quantifies the amount of “noise” in changes in allele/trait frequencies due to finite population size *N*, i.e., genetic drift.

The drift vector describes the expected change in mutant allele frequencies in time due to selection, mutation, and recombination. For the trait frequencies, this term is especially complex.

Using (3) and (4) and dropping infinitesimals smaller than 𝒪 (1*/N*), we find

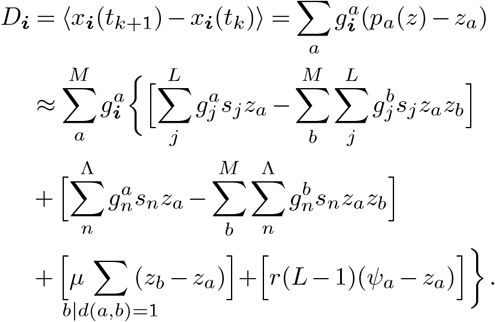

Here we have four terms, which describe the evolutionary forces of selection for individual mutant alleles, selection on binary traits, mutation, and recombination, respectively.

Contributions due to natural selection can be written in the same way for both mutant alleles *i* and traits *n* (which, as a reminder, we group under a general bold-faced index ***i***). Using (6) and the expression for pair allele/trait frequencies 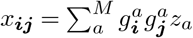, these terms become

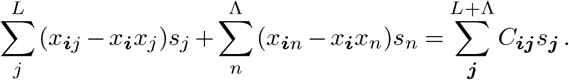

The mutation and recombination terms are different for mutant alleles and for traits. Here, we start with the mutation term for mutant alleles, which can be simplified as

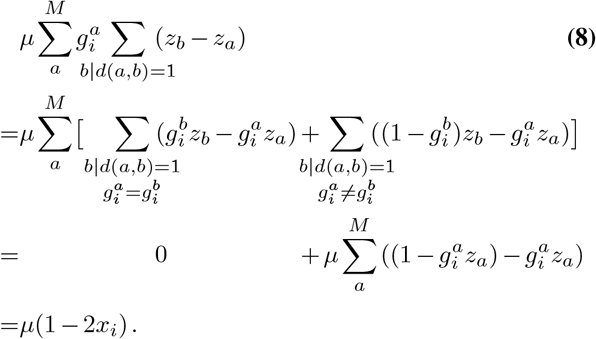

For the first term, *b*|*d*(*a, b*) = 1, 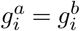 means that *a* and *b* have the same allele at locus *i*, but they differ by one mutation at some other locus. If we sum over every genotype, these terms make no contribution to the change in mutant allele frequency *i* because every pair (*a, b*) has one conjugate pair (*b, a*). The second term is not symmetrical, but every *a* only has one *b* that can satisfy the conditions *b*|*d*(*a, b*) = 1, 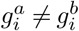(i.e., the genotype with a MT allele at site *i* flipped to WT, or vice versa). Thus, we can use *a* to replace *b* and its summation.

Physically, (8) expresses the total flux due to mutation, which is the probability of mutation that can change the allele at locus *i* from WT to MT (flux in) minus the probability from MT to WT (flux out). The probability of MT at locus *i* is the mutant allele frequency. Since the state for the allele is binary, the sum of MT frequency and WT frequency is 1. Thus, we can easily simplify the expression as

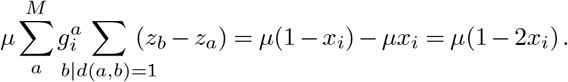

Next, we will consider the recombination term for mutant alleles. Similar to the mutation term, this represents the recombination flux for locus *i*. However, the expected mutant allele frequency change due to recombination alone is always zero. This is because, for every case in which a sequence without a mutant allele recombines with a sequence that has a mutant allele such that the recombinant sequence has the mutant allele (thus increasing the mutant allele frequency), there is another case with the MT and WT sequences switched (thus decreasing mutant allele frequency by the same amount) which occurs with the same probability. Thus, this term vanishes by symmetry.

In total, then, the drift vector for mutant frequencies *i* is

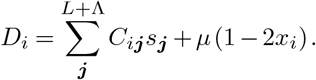

Now we consider the mutation and recombination terms for traits. These terms are more complex because the conditions that can change the trait are different from the ones for a single locus, and generally depend on the state of all alleles that contribute to the trait. For example, mutation at one site within a trait group *n* will change the allele from WT to MT (or vice versa), but it may not change the state for the trait as a whole if there are other mutations among the trait group.

Let us consider the mutation term for traits, which we can expand as

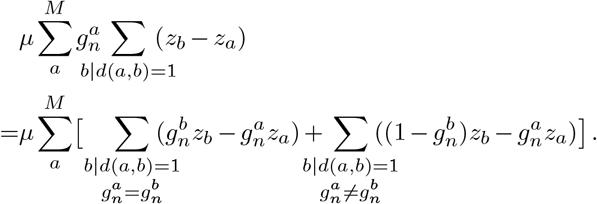

As in (8), the first term is also zero. However, the second mutation term is different. In this case, genotypes *a* and *b* that satisfy *b*|*d*(*a, b*) = 1, 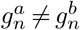 do not exhibit a one-to-one correspondence: if all alleles in the trait group for *a* are WT, for example, then there are many *b* can satisfy these conditions and the number of *b* is the length of the trait group *n*. Alternately, if genotype *a* contains more than one mutant allele in the trait group, then it cannot be changed to WT within a single mutation.

To address this issue, we introduce a new variable, denoted as 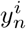, which is the frequency of genotypes that contain only one mutation in trait group *n*, and the mutation is at locus *i*. This can be written as

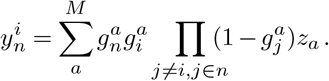

With this new variable, we can rewrite the mutation term for traits. Physically, to change the trait value (starting from WT), every locus among the trait group needs to be considered, as any mutation in these loci can change the state of the trait group from WT to MT. Conversely, mutation can only change the trait value from MT to WT if it affects genotypes that have one mutation among the trait group (i.e., by converting the single mutant allele to WT). Thus,

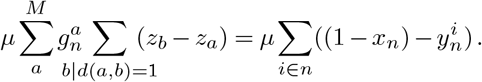

Finally, we turn to the recombination term for traits. Here we start by building physical intuition for the result. In each generation of the evolutionary process, recombination occurs numerous times. For this issue, we need to focus solely on the recombination that can alter the state of a trait group. In particular, we must enumerate the probabilities of recombination events that change traits *and* for which the reverse process (i.e., switching the order of the recombination partner sequences, which has the same probability) does not change the trait in the opposite way, thus yielding no net expected change in the trait frequency.

We use the index *k* to represent a recombination break-point. To change a trait’s value, *k* must be located after the first site in the trait group and before the last. Since we have assumed that the recombination probability per site per generation *r* is small, we will only consider cases where a single recombination breakpoint falls within the trait sites, but the analysis below could be generalized to allow for two or more breakpoints. If recombination occurs between two sequences, where one is WT for trait group *n* and the other is MT before and after a breakpoint *k* for the same trait, then all the recombinants will be MT for trait group *n*. For such sequences, recombination can change WT to MT for the trait, but not MT to WT. This is the only scenario in which recombination leads to a net *increase* in trait frequency.

In the opposite direction, consider recombination between two sequences, where one is WT for trait *n* before a breakpoint *k* and MT after, and the second sequence is MT before *k* and WT after. Thus, both sequences are MT for trait *n*, but through recombination a WT sequence can be produced. This is the only recombination process that leads to a net *decrease* in trait frequency.

Collecting these two terms together, we arrive at the following expression for the net expected change in trait frequency due to recombination,

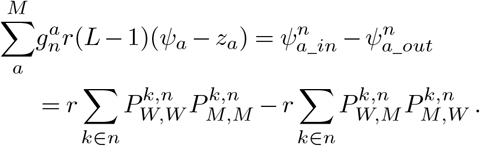

Here 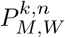 represents the frequency of sequences that have at least one mutant allele in the trait group *n* on or before site *k*, and all WT alleles in the same trait group after *k*. In total, then, the drift vector for traits is

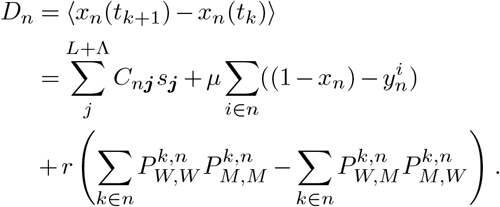

### Two-step model of HIV-1 recombination

As noted in the main text, HIV-1 recombination actually occurs in multiple steps, including the coinfection of a single cell by two distinct viruses and reverse transcriptase template switching between the two distinct RNA strands. Let us refer to the probability of coinfection as *p*_*c*_ and the probability of template switching as *p*_*s*_ per base per replication cycle. We explored how this two-step model of recombination would affect our estimates of selection.

As before, we must search for recombination events that could affect the escape frequency for a particular epitope. In this model as well, recombination breakpoints must occur within an epitope (i.e., after the first site in the epitope and before the last site) to affect escape frequency. Previously, we had assumed a single effective recombination rate per site *r* ≪ 1. Now, we break this into two steps, with probability *p*_*c*_ for coinfection and *p*_*s*_ per base for template switching. Naively relating *r* ∼ *p*_*c*_*p*_*s*_, the difference between these models becomes clear: the probability of observing two recombination breakpoints within some region is proportional to *r*^2^ in the one-step model, but 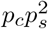 in the two-step model.

Previously, we had assumed that multiple recombination events within the same epitope should be very unlikely, and that the dominant contribution to changes in escape frequency should therefore come from single recombination events (i.e., *r*_2_ ≪ *r*). To make the same approximation in the two-step recombination model, we would need to show that 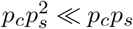.While estimates for the template switching probability vary, high-end estimates are generally around *p*_*s*_ ∼ 10^−3^. There is also evidence for potential hotspots of recombination in HIV-1, where template switching may occur more frequently ^52–55^. However, even if template switching were to occur at a rate of 10^−2^ per base in a recombination hot spot, the assumption that 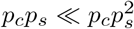 is likely to be a very reasonable one. Thus, to leading order in the template switching rate, one could replace the more biologically realistic two-step recombination process with an effective one-step process with an effective rate *r* = *p*_*c*_*p*_*s*_ to extract the dominant contributions of recombination to escape frequency change (and therefore to inferred selection).

### Marginal path likelihood (MPL) inference

To find the fitness effects of alleles/traits that best fit the data, we will attempt to find parameters *s*_***i***_ that maximize the likelihood of the data. Since the likelihood *P* is proportional to action ***S*** (see (7)), the unknown selection coefficients *s*_*i*_ and trait coefficients *s*_*n*_ that can maximize action ***S*** will maximize the path probability *P*.

To control our estimates, we also introduce a Gaussian prior distribution for the selection coefficients with mean zero and precision *γN*,

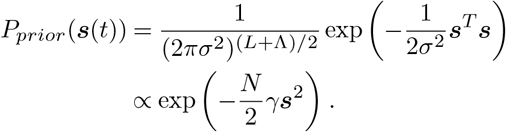

Here we have absorbed the population size *N* into the width of the prior for convenience (see below). Including the prior distribution, the action becomes

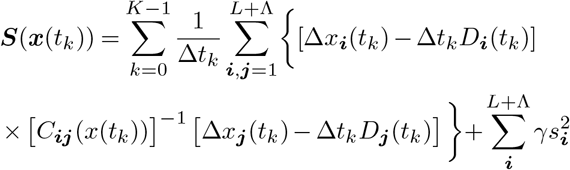

Finally, we can apply Bayes’ theorem to infer selection coefficients *ŝ*_***i***_ that maximize the posterior distribution, providing the best compromise between the prior and the likelihood. While the posterior distribution is a complicated function of allele/trait frequencies, it is simply a Gaussian function of the selection coefficients. This allows us to write an analytical expression for the maximum a posteriori parameters, as shown in equation (2) in the main text, with

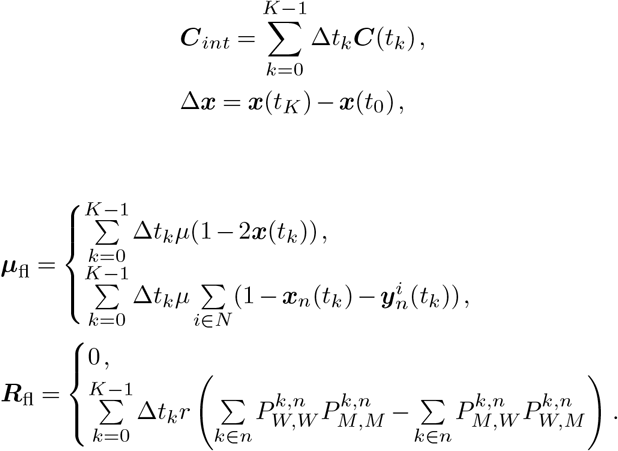

Here, the top expressions for ***µ***_fl_ and ***R***_fl_ are for mutant alleles, and the bottom ones are for traits. In simulations, we used *γ* = 1 for selection coefficients and 0.1 for traits. In noisier HIV-1 data, we used *γ* = 10 for selection coefficients and 1 for traits.

### Extension to multiple alleles per locus and asymmetric mutation probabilities

To study real sequences, we can extend the simple model presented in the previous sections to allow for multiple alleles per locus and asymmetric mutation probabilities. We use *α, β*,… indices to represent different alleles (ranging, for example, over nucleotides or amino acids), where we write the total number of possible alleles at a locus as *l*. Our fitness model is then:

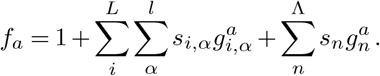

Similarly, *s*_*i*_,_α_ represents the selection coefficient for allele *α*at locus *i*, and 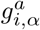 equals 1 if genotype a has allele *α* at locus *i*. T he trait term is the same as the binary case since it does not have a natural counterpart (only 2 states, WT and MT). We use *x*_*i,α*_(*t*_*k*_) to represent the frequency of allele *α* at locus *i* at generation *t*_*k*_, and *µ*_*αβ*_ to denote the probability per locus per generation of mutation from allele *α* to *β*. Following parallel arguments to before, the MPL estimate of the selection coefficient *s*_*i,α*_ for each allele *α* at each locus *i* and the trait coefficients *s*_*n*_ can be obtained.

First, we write the diffusion matrix *C*_*iα,jβ*_(*t*_*k*_)*/N*,

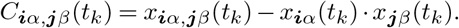

where *x*_***i****α*,***j****β*_(*t*_*k*_) is the frequency of sequences with alleles *α* and *β* at loci ***i*** and ***j*** at generation *t*_*k*_. When one of the indices corresponds to a trait, allele subscripts are not needed. For example, the covariance between trait group *n* and allele *α* at locus *i* can be written as *C*_*n,iα*_(*t*_*k*_) = *x*_*n,iα*_(*t*_*k*_) − *x*_*n*_(*t*_*k*_) · *x*_*iα*_(*t*_*k*_).

The estimates selection coefficients for alleles are then

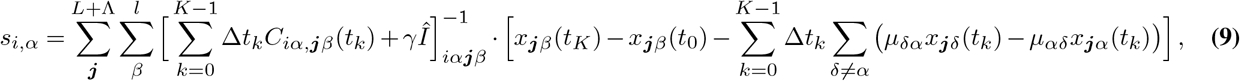

And the selection coefficients for traits are given by

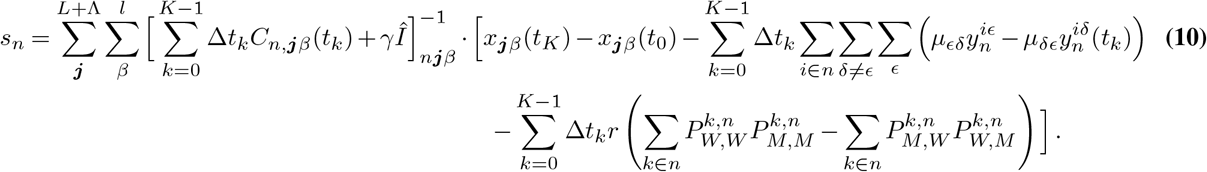

Here the index *ϵ* runs over WT alleles and synonymous mutations for each binary trait *n*. This is because only nonsynonymous mutations contribute to changes in the trait in our model.

In this version of the model, we compute selection coefficients for both mutant and WT alleles. However, what is important is not the selection coefficient for an individual allele in isolation, but rather the change in fitness upon mutation. Thus, the selection coefficients we report are normalized by subtracting the WT selection coefficient from the selection coefficient of each allele at each locus (*s*_*iα*_ − *s*_*iW*_). In this way, the selection coefficients are normalized such that the WT selection coefficient is 0, and the selection coefficients for other alleles quantify the advantage or disadvantage relative to WT. In the limit that the selection coefficients are small, one can show that the WF dynamics are invariant under such shifts in the selection coefficients. In physics, this phenomenon is referred to as gauge invariance.

### Simulation data

We simulated the WF model with discrete generations and binary (mutant/WT) states in Python. Briefly, we evolved populations of sequences according to (5) over multiple generations, starting with an initial population of all WT sequences and recording the entire evolutionary history. After simulation, we computed the single *x*_*i*_, double *x*_*ij*_ mutant frequencies, and trait frequencies *x*_*n*_ from the sampled sequences trajectories and used them to infer individual selection coefficients (9) and trait coefficients (10). Parameter values are detailed in **Fig. 1**. The simulation and analysis code with original simulation data are contained in the GitHub repository.

Since real data typically contains only a small portion of population, and is not sampled at every generation, we also studied how different sampling depths and sampling time intervals affect the performance of our method. We chose part of the sequences from the population and time points to estimate individual selection and trait coefficients. *n*_*s*_ denotes the number of sequences we randomly selected from the population and Δ*t* is the time interval, which means we choose the data every Δ*t* generations. The initial population and simulation parameters are described in **Supplementary Fig. 2**.

CD8^+^ T cell levels can fluctuate over time, suggesting that a time-varying binary trait selection coefficient would most accurately describe this phenomenon. Thus, we also conducted a simulation with time-varying selection coefficients of binary traits to test our approach. The results in **Supplementary Fig. 1** demonstrate that the trait coefficients we estimate are approximately equivalent to the average trait coefficient when selection varies over time. The parameter values are detailed in **Supplementary Fig. 1**.

### HIV-1 sequence data

We obtained HIV-1 sequence data from 13 individuals of the CHAVI 001 and CAPRISA 002 studies in the United States, Malawi, and South Africa from the Los Alamos National Laboratory (LANL) HIV Sequence Database. We applied several selection criteria ^50^ to minimize the influence of noise in the data, including removing the sequences with large numbers of gaps, sites with high gap frequencies, and time points with very small numbers of sequences or large gaps in time from the last sample. We also imputed ambiguous nucleotides with the most common nucleotides observed at the same site within the same individual.

For these 13 individuals, sequence data consisted of 3^*′*^ and 5^*′*^ half-genome sequences, which were approximately 4,500 bp in length. Our analysis focused only on polymorphic sites, where more than one nucleotide (including gaps/deletions) was observed in an individual (approximately 100-900 bp in length). To infer selection, we used a mutation rate matrix estimated in ref. ^56^ as input. We allowed the effective recombination rate *r* to vary along with patient viral load (VL), following recent work that revealed increasing effective recombination rates in individuals with higher VL due to higher levels of coinfection ^57^. To estimate *r* as a function of VL, we used a linear model with parameters roughly fit to the data of Romero and Feder (*r* = 1.722 × 10^−10^ VL + 1.39 × 10^−5^).

In our model, we dynamically determined the recombination rate based on the viral load at each time point, which was measured in past work ^58^. For later stages of infection where VL was not measured, we assumed that VL values remained unchanged from the most recent measurement, consistent with the establishment of a viral set point in chronic infection. Although there are large spikes in VL during acute infection (resulting in a corresponding recombination rate on the order of 10^−3^, compared to typical constant estimates of around 1.4 × 10^−5^ (ref. ^59^)), it quickly settles down to a value orders of magnitude lower. We treated the transmitted/founder (TF) sequence as the “wild-type” sequence for each individual.

The locations of CD8^+^ T cell epitopes for these sequences were experimentally ^58^ or computationally ^60^ determined. *Escape sites* refer to polymorphic sites where nonsynonymous mutations were observed in the reading frame of a CD8^+^ T cell epitope. In this way, we anticipate that nonsynonymous mutations in escape sites will affect T cell recognition. We consider the escape sites that can change the same epitope are among one trait group.

### Calculation of effects of linkage on inferred selection

Δ*ŝ*_*ij*_ can tell us the effects of linkage from variant *i* to *j*. To compute it, we calculate the coefficients for mutant variant *j* and eliminate the influence from mutant variant *i* by artificially reverting variant *i* to WT ^50^.

For mutant alleles, we generated a modified version of the sequence data where all mutant variants *i* are replaced by the corresponding TF nucleotide for all sequences at all time points. For traits, we treat all mutant variants within one epitope as WT. With these modified data, we can infer the coefficients again for all variants ***j***, denoted as 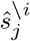.Then we define

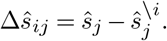

Positive values of Δ*ŝ*_*ij*_ indicate that linkage with variant *i* can increase the selection coefficient inferred for variant *j* and vice versa. By computing the Δ*ŝ*_*ij*_, we can quantify the effect of linkage on inferred selection.

### The effects of recombination on the inference

Unlike selection on individual alleles, which has been studied previously using similar approaches ^50,51^, the selection that we infer for binary traits is affected by recombination. How large is the contribution of recombination in this analysis?

To answer this question, we inferred selection on mutant alleles and traits in simulations and in the HIV-1 data sets, with and without the inclusion of recombination. In general, we find that the effect of recombination on the inferred coefficients is small. This is reasonable, as recombination only affects the traits, which constitute a small portion of a sequence. For simulation data, there are three traits in a sequence length of 50; however, for experimental data, it is often a few epitopes contained within sequences that are thousands of base pairs long.

Simulation results (**Supplementary Fig. 7**) illustrate that the recombination term has negligible effects on the inference of individual selection coefficients. For trait terms, incorporating recombination does lead to small but noticeable improvements in inference. In HIV-1 data, the influence of the recombination term is small (**Supplementary Fig. 8**). Although the recombination rate in HIV-1 is relatively high, the scenarios under which recombination will lead to net change in trait frequencies are rare.

## Data and code

Data and code used in our analysis is available in the GitHub repository located at https://github.com/bartonlab/paper-binary-trait-inference.

This repository also contains Jupyter notebooks that can be run to reproduce our figures and analysis.

---

**Supplementary Fig. 1.**
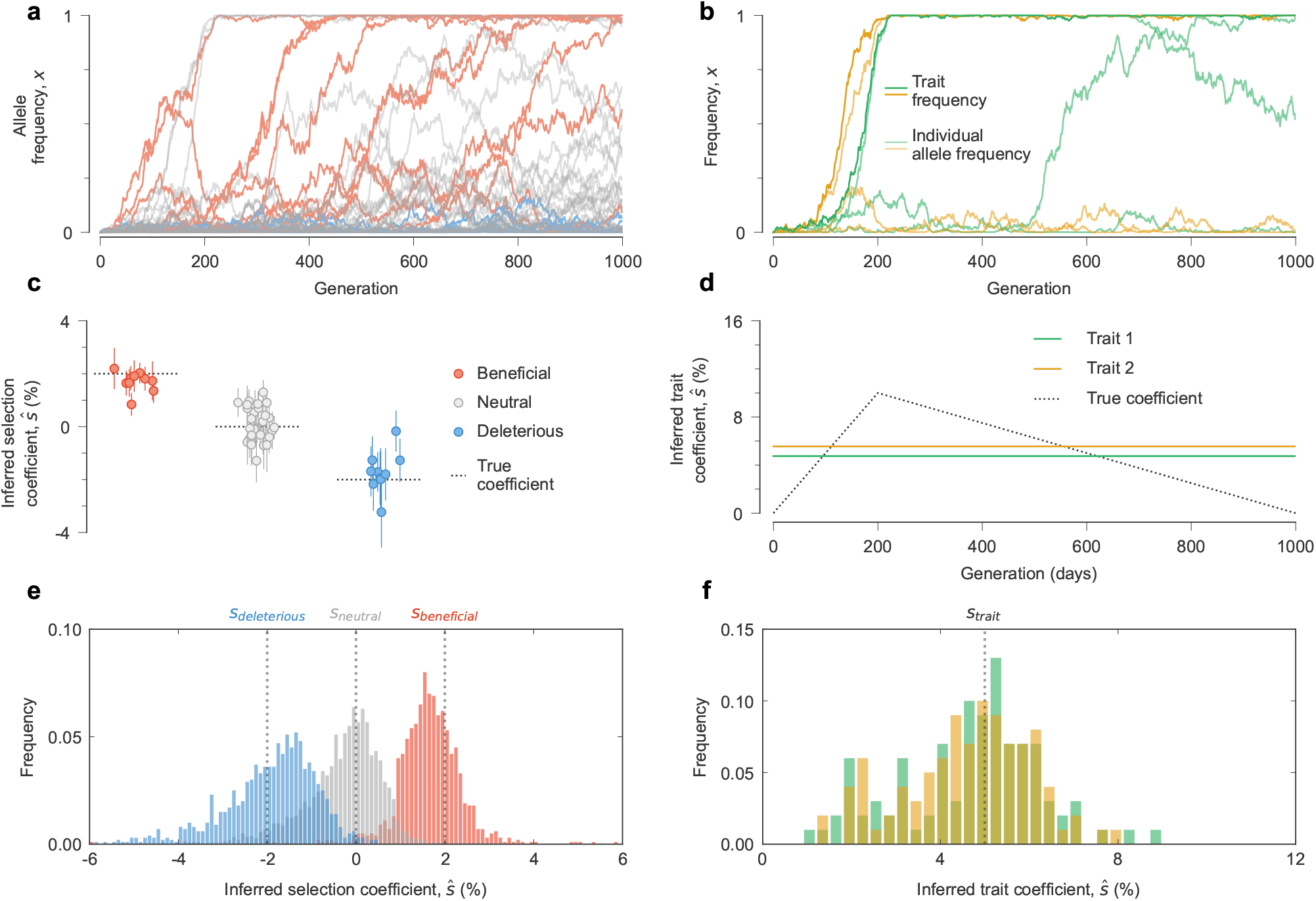
MPL recovers selection from data with time-varying selection coefficients for binary traits. **a**, Simulated mutant allele frequency trajectories. **b**, Trait frequencies and their contributing individual mutant allele frequencies in the same simulation. The fitness contributions of individual mutations (**c**) and traits (**d**) that we infer are close to their true values. The true values for the selection coefficient for individual loci are constant while those for binary traits vary over time. Inferred trait coefficients are close to the average values of time-varying coefficients over the course of the simulation. Distribution of (**e**) individual selection coefficient and (**f**) trait coefficient estimates across 100 replicate simulations. The true value for the selection coefficients for the binary traits shown here is the average value. Simulation parameters: *L* = 50 loci with two alleles at each locus (mutant and WT), ten beneficial mutants with *s* = 0.02, 30 neutral mutants with *s* = 0 and ten deleterious mutants with *s* = −0.02. We consider two binary traits, each with three contributing alleles and time-varying trait coefficients. The selection coefficients for these two binary traits go from 0 to 0.1 over the beginning 200 generations and then go down to 0 over the remaining 800 generations. Mutation probability per site per generation *µ* = 2 *×* 10^−4^, recombination probability per site per generation *r* = 2 *×* 10^−4^, population size *N* = 10^3^. The initial population contains all WT sequences, evolved over *T* = 1000 generations.

**Supplementary Fig. 2.**
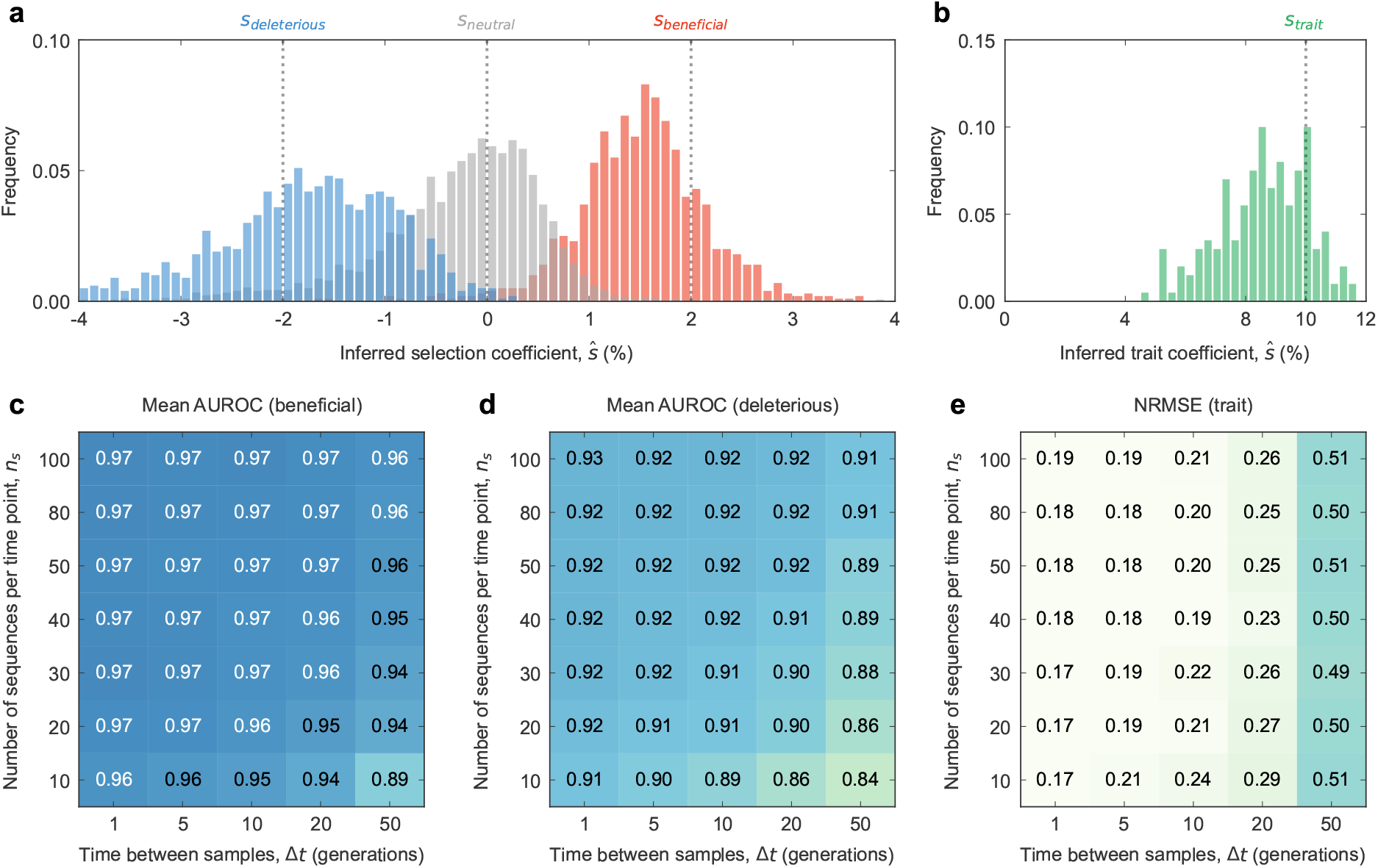
MPL recovers selection from complex dynamics even from limited data. Distribution of (**a**) individual selection coefficient and (**b**) trait coefficient estimates across 100 replicate simulations, using the same parameters as in **Fig. 1**. Our approach is robust to finite sampling constraints, as measured by the accurate classification of (**c**) beneficial and (**d**) deleterious mutants and (**e**) inference of trait coefficients, even when the number of sequences sampled per time point *n*_*s*_ is low and the spacing between time samples Δ*t* is large. AUROC, area under the receiver operating characteristic; NRMSE, normalized root mean square error. Simulation parameters: *L* = 50 loci with two alleles at each locus (mutant and WT), ten beneficial mutants with *s* = 0.02, 30 neutral mutants with *s* = 0, and ten deleterious mutants with *s* = −0.02. We consider Λ = 2 trait groups, each with three contributing alleles and trait coefficients *s* = 0.1. Mutation probability per locus per generation *µ* = 2 *×* 10^−4^, population size *N* = 10^3^. The initial population is all wild type, evolved over *T* = 1000 generations.

**Supplementary Fig. 3.**
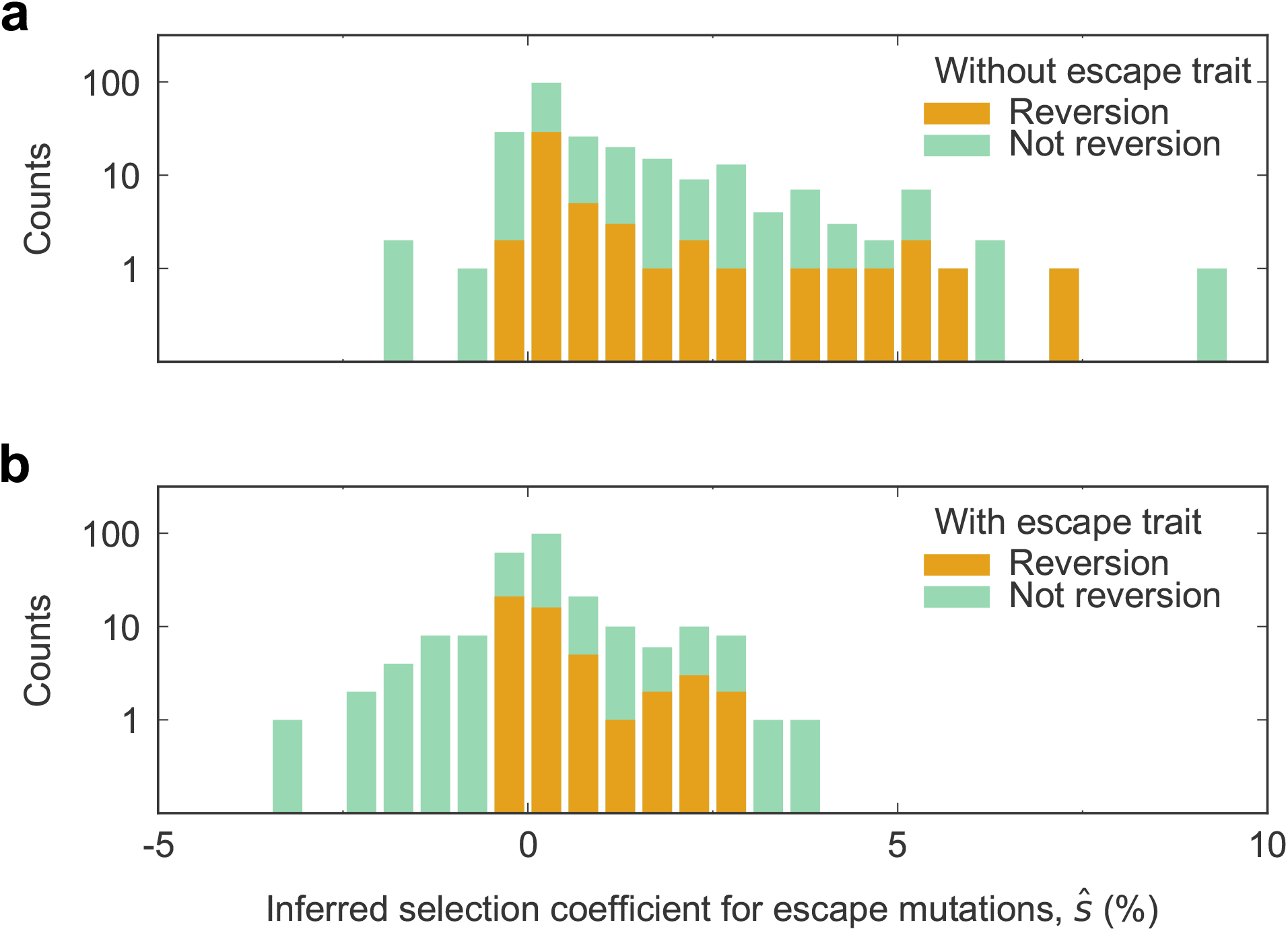
Inclusion of escape coefficients substantially shifts inferred selection coefficients for escape mutations toward more deleterious values. **a**, Distribution of inferred selection coefficients without escape traits. **b**, Distribution of inferred selection coefficients with escape traits. Reversions are slightly more likely to be inferred to be beneficial/less like to be deleterious than non-reversions.

**Supplementary Fig. 4.**
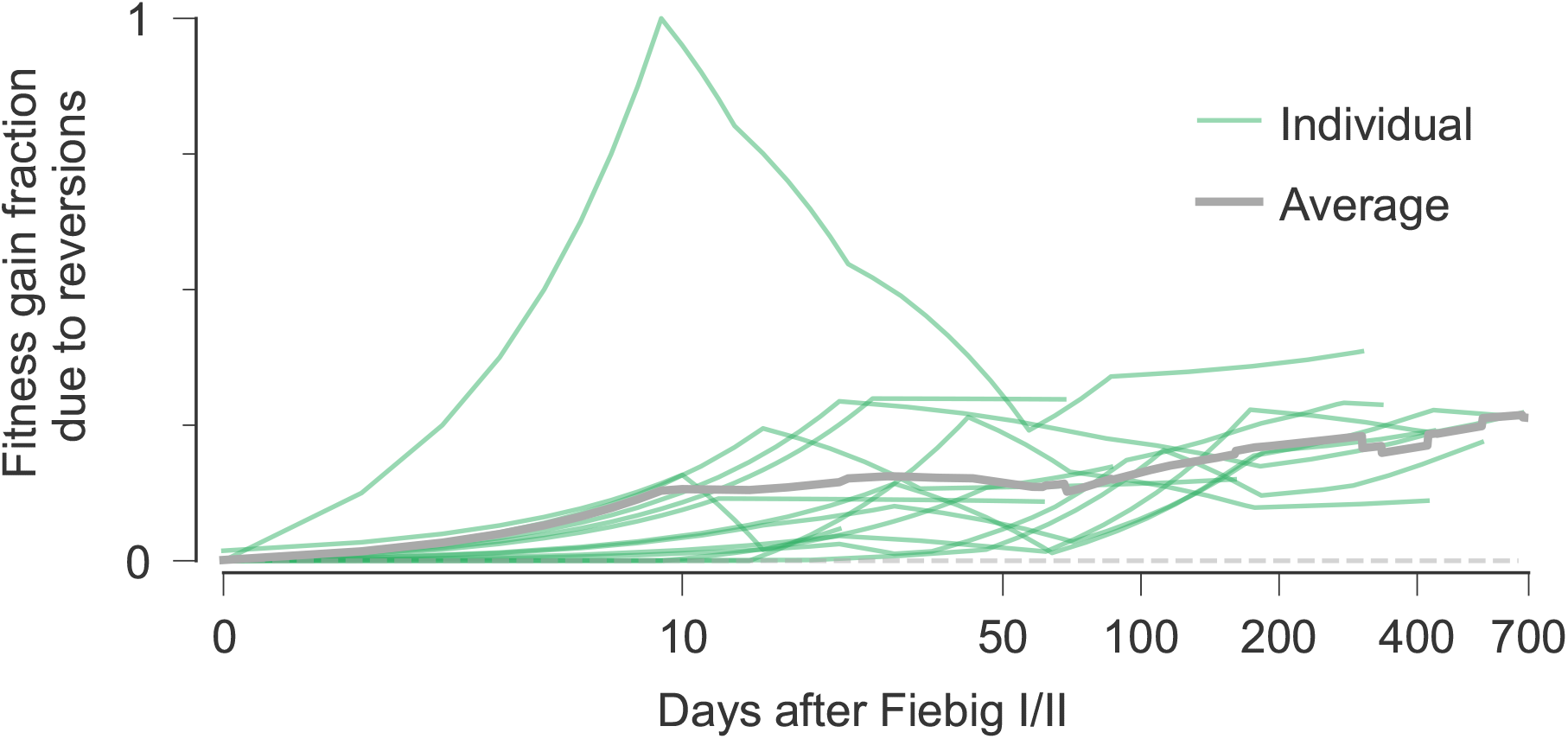
The contribution of reversions to intrahost HIV-1 fitness gains is significant and grows over time. Reversions refer to mutations that revert from the transmitted/founder (TF) variant to the nucleotide of the HIV-1 clade consensus sequence.

**Supplementary Fig. 5.**
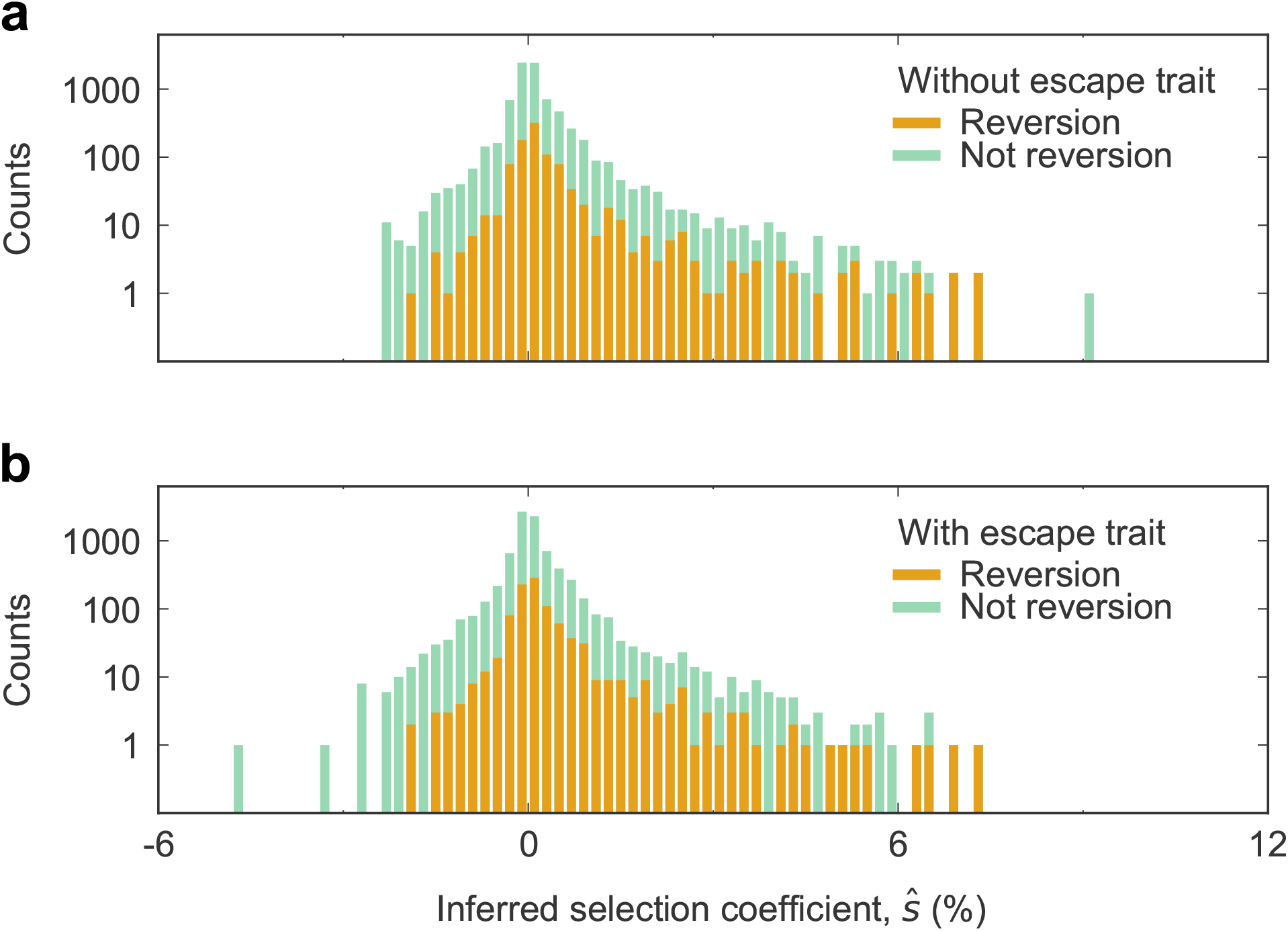
The distribution of inferred selection coefficients with and without the inclusion of escape traits. **a**, Distribution of inferred selection coefficients without escape traits. **b**, Distribution of inferred selection coefficients with escape traits. We note an enrichment in substantially beneficial mutations that are also reversions to the HIV-1 clade consensus sequence. While some large-effect mutations shift substantially with the inclusion of escape traits (i.e., escape mutations), the effect on the bulk of the inferred selection coefficients is minimal.

**Supplementary Fig. 6.**
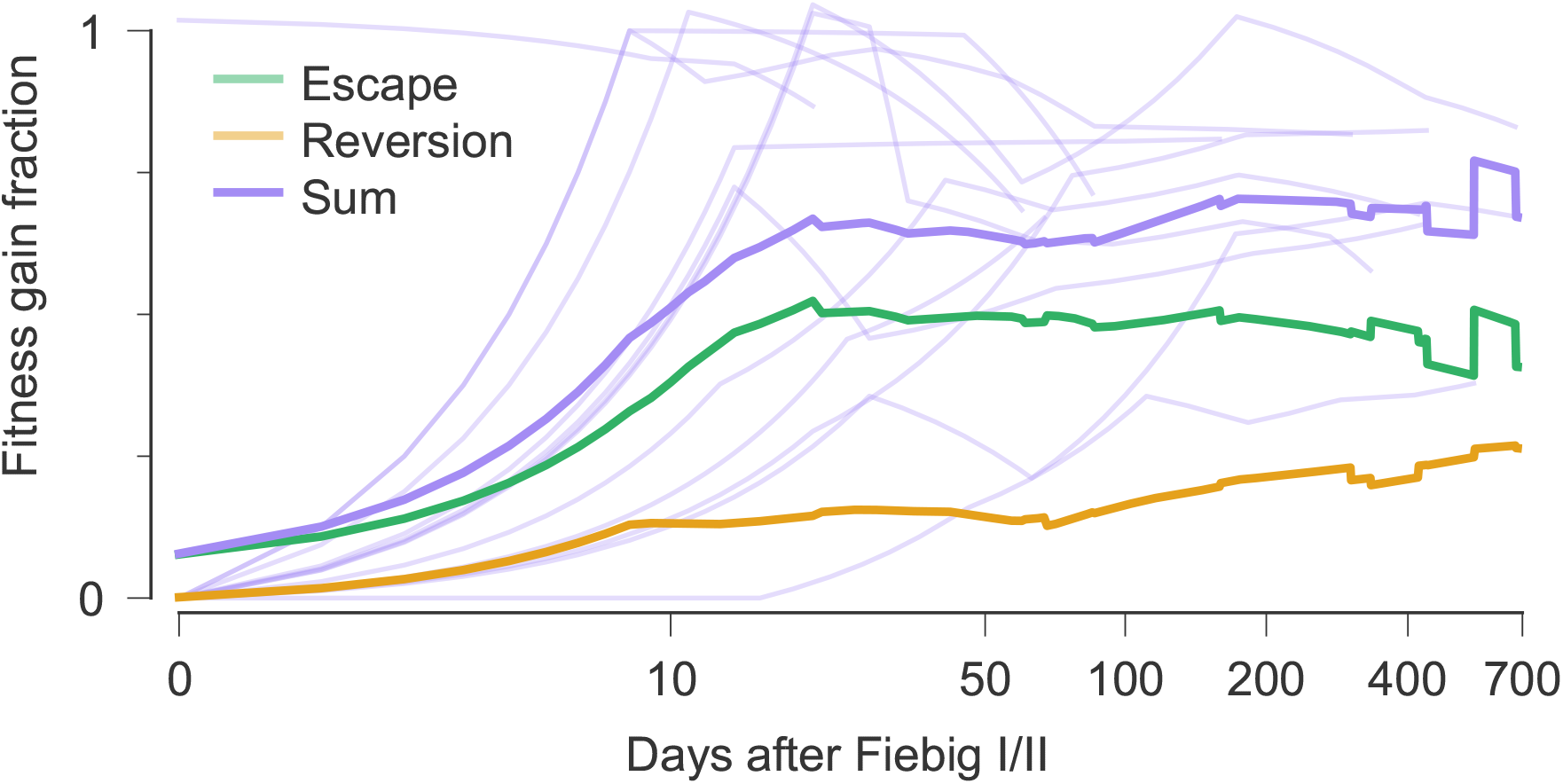
Intrahost HIV-1 fitness gains due to CD8^+^ T cell escape mutations and reversions. In total, CD8+ T cell escape (see **Fig. 2b**) and reversions (see **Supplementary Fig. 4**) make dominant contributions to HIV-1 fitness gains *in vivo*. Across the 13 patient data sets that we studied, the fraction of fitness gains due to escape and reversions is around 75% on average. At earlier times, T cell escape plays a larger role, while the contribution of reversions grows steadily over time.

**Supplementary Fig. 7.**
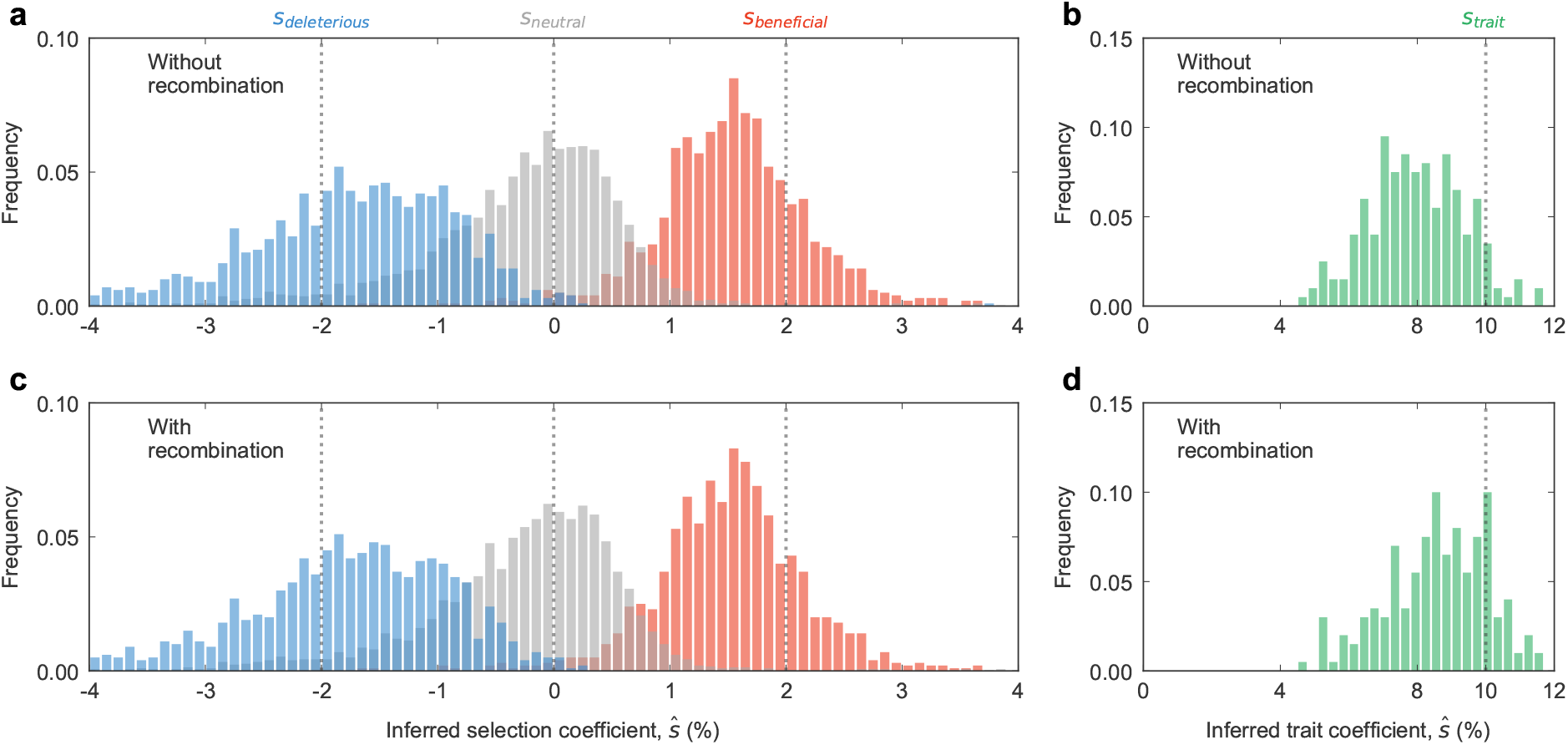
Effects of recombination on inferred selection coefficients and trait coefficients in 100 replicate simulations. The distribution of inferred (**a**) selection coefficients and (**b**) trait coefficients in simulations, using the same parameters as in **Fig. 1**, without including the effects of recombination in the estimator. **c, d**, Analogous distributions of inferred coefficients when recombination is included in the estimator. The effects of recombination on the inferred parameters are subtle, with the most prominent feature being a shift in the inferred trait coefficients toward the true value and away from zero.

**Supplementary Fig. 8.**
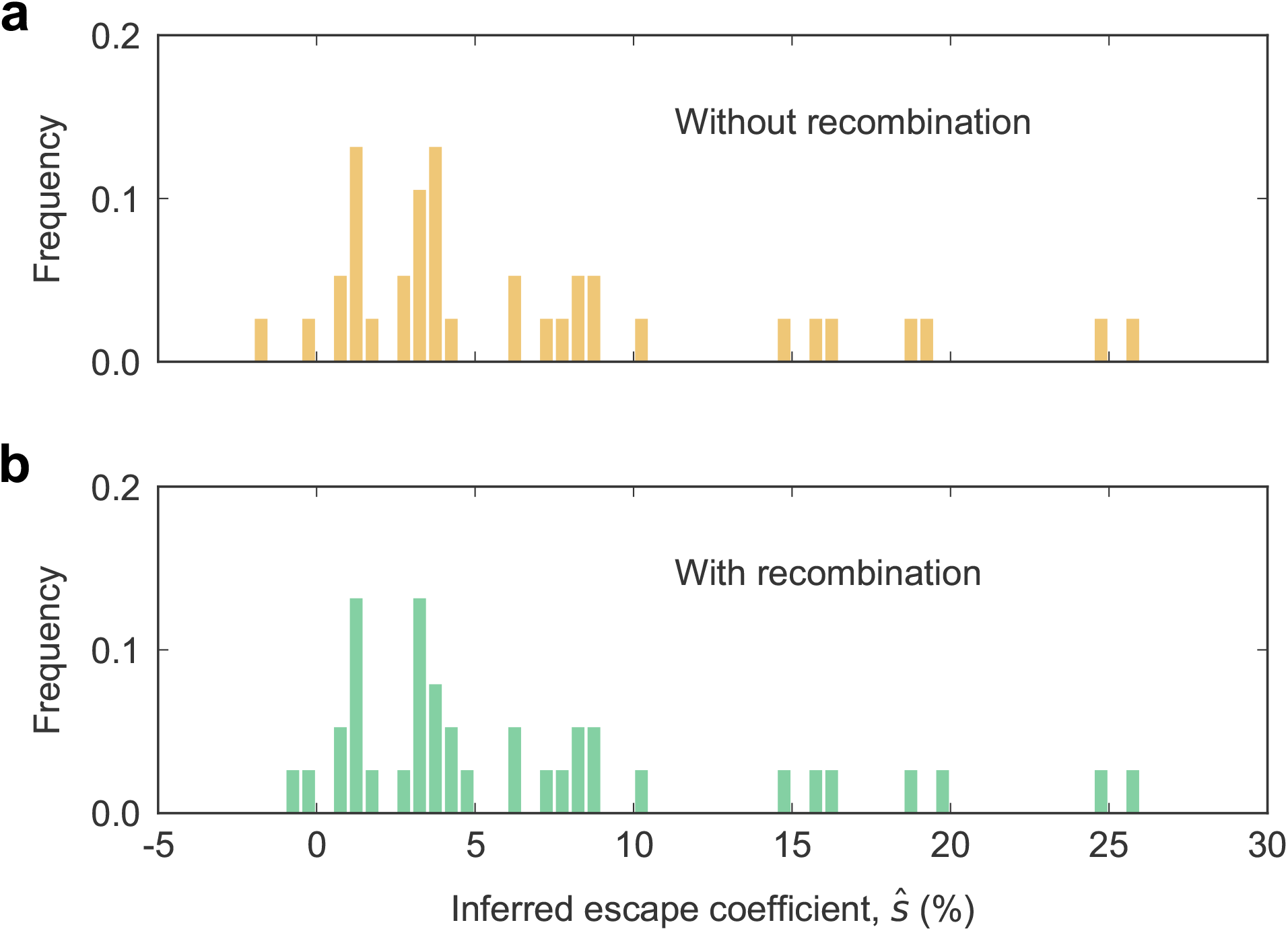
Effects of recombination on inferred escape coefficients in HIV-1 data. Distribution of inferred escape coefficients (**a**) without and (**b**) with the inclusion of recombination in the estimator. Even though the recombination rate for HIV-1 is relatively high (*r* ∼ 1.4 *×* 10^−5^), recombination events capable of altering trait frequencies are rare. The contribution of ***R***_fl_ to the inferred escape coefficients is thus small.

